# The contribution of rare variation to prostate cancer heritability

**DOI:** 10.1101/023440

**Authors:** Nicholas Mancuso, Nadin Rohland, Kristin Rand, Arti Tandon, Alexander Allen, Dominique Quinque, Swapan Mallick, Heng Li, Alex Stram, Xin Sheng, Zsofia Kote-Jarai, Douglas F. Easton, Rosalind A. Eeles, the PRACTICAL consortium, Loic Le Marchand, Alex Lubwama, Daniel Stram, Stephen Watya, David V Conti, Brian Henderson, Christopher Haiman, Bogdan Pasaniuc, David Reich

## Abstract

Although genome-wide association studies (GWAS) have found more than a hundred common susceptibility alleles for prostate cancer, the GWAS reported variants jointly explain only 33% of risk to siblings, leaving the majority of the familial risk unexplained. We use targeted sequencing of 63 known GWAS risk regions in 9,237 men from four ancestries (African, Latino, Japanese, and European) to explore the role of low-frequency variation in risk for prostate cancer. We find that the sequenced variants explain significantly more of the variance in the trait than the known GWAS variants, thus showing that part of the missing familial risk lies in poorly tagged causal variants at known risk regions. We report evidence for genetic heterogeneity in SNP effect sizes across different ancestries. We also partition heritability by minor allele frequency (MAF) spectrum using variance components methods, and find that a large fraction of heritability (0.12, s.e. 0.05; 95% CI [0.03, 0.21]) is explained by rare variants (MAF<0.01) in men of African ancestry. We use the heritability attributable to rare variants to estimate the coupling between selection and allelic effects at 0.48 (95% CI of [0.19, 0.78]) under the Eyre-Walker model. These results imply that natural selection has driven down the frequency of many prostate cancer risk alleles over evolutionary history. Overall our results show that a substantial fraction of the risk for prostate cancer in men of African ancestry lies in rare variants at known risk loci and suggests that rare variants make a significant contribution to heritability of common traits.

## Introduction

More than 220,000 men are expected to be diagnosed with prostate cancer (PrCa) and more than 27,000 are expected to die of the disease in the United States alone in 2015^1^. Approximately 58% of risk for PrCa is estimated to be due to genetic inherited factors^2-6^. To date, genome-wide association studies (GWAS) have revealed more than 100 common risk variants for PrCa that explain ~33% of the familial risk^7-25^ leaving the majority of risk unexplained (the “missing heritability” problem). Since GWAS have primarily investigated common variants (MAF>1-5%) for association to PrCa risk, an unexplored hypothesis is that part of the “missing heritability” of PrCa risk is attributable to rare (MAF<0.01) variants. To address this hypothesis we focused on examining rare variation at known susceptibility regions that are only partially tagged by GWAS arrays. The rationale for investigating known risk regions is that a) unlike the rest of the genome, genetic variation in these regions has been established to confer risk and b) there are examples of rare and low-frequency variation at known GWAS-identified risk regions being important for a number common diseases, including prostate cancer (i.e. 8q24)^26-29^.

We conducted targeted sequencing of known PrCa GWAS loci to investigate the contribution of low frequency and rare variation to PrCa risk. We targeted all 63 autosomal PrCa risk regions that were known to us at the time of study design (since then, an additional 29 loci have been discovered and we did not interrogate them for this study). For each region, we started with the index SNP previously associated to PrCa by GWAS, and then attempted to tile Agilent SureSelect baits covering all nucleotides within a block of strong linkage disequilibrium around the SNP (plus exons and conserved elements within 200kb of the SNP). We made individually barcoded next-generation sequencing libraries for all of the samples, pooled these libraries in sets of typical size 24, and then performed an in-solution hybrid enrichment. After removal of duplicated molecules, we achieved an average depth of 9.3x coverage in 9,237 cases and controls across four ancestry groups (4,006 African, 1,753 European, 1,770 Japanese and 1,708 Latino). We identified 197,786 variants across all ancestries, imputed genotypes in all individuals, and then correlated them to PrCa risk.

First, our analysis shows that sequencing of a large set of individuals interrogates substantially more of the genetic variance for the trait than studies of GWAS variants alone, as the variance explained in the trait by all the sequenced variants is significantly larger than variance explained by known GWAS variants at the same loci. Second, we find evidence of genetic heterogeneity by ancestry in risk for prostate cancer. Third, we use variance component methods to partition the SNP-heritability across different frequency classes and find a large amount of SNP-heritability coming from the rare variant class in men of African ancestry (e.g. variants with 0.1%≤MAF<1% explained 0.12 of variance in the trait as compared to 0.17 for MAF≥1% variants). We used the SNP-heritability assigned to the rare variant class to make the first relatively precise estimate of the strength of coupling between selection and allelic effect for a common trait. Finally, we replicated association signals at known GWAS loci and utilized an approach that combines epigenetic annotation data (e.g. localization of androgen receptor sites in a prostate adenocarcinoma cell line) with the association signal to identify plausible causal variants at some of these loci.

## Results

### Experimental strategy

To explore the contribution of rare and low frequency variation to risk of PrCa, we targeted 90 index SNPs at 63 autosomal risk regions that had been associated to PrCa risk by GWAS at the time that this study was designed (October 2011). For each index SNP, we used Haploview^30^ (HapMap release 24) to visualize the surrounding block of linkage disequilibrium (LD) in European ancestry individuals. We then manually identified boundaries for target capture based on the region where |D′| values fell precipitously (see Supplementary Figure 1 and Supplementary Note 2). We also targeted all exons (defined based on RefSeq) within 200kb of each index SNP and all conserved non-coding sequences (defined based on the 29 mammals project^31^) within 5kb of each exon, and elements > 100bp in length or with conservation scores >75 within the 200kb window around each index SNP. Outside of the targeted GWAS loci, we also included exons and conserved elements of *MYC* and *PVT1* because of their potential importance to PrCa. We designed and ordered Agilent SureSelect^32^ in-solution enrichment probes to target a total of 12 Mb in the regions after two rounds of target design which was restricted to 11.1Mb in the analyses after QC. The total span of regions we wished to target was 16.7 Mb, but we were not able to design probes for 4.7 Mb due to repetitive elements that needed to be masked during probe design (Supplementary Note 2).

**Figure 1.**
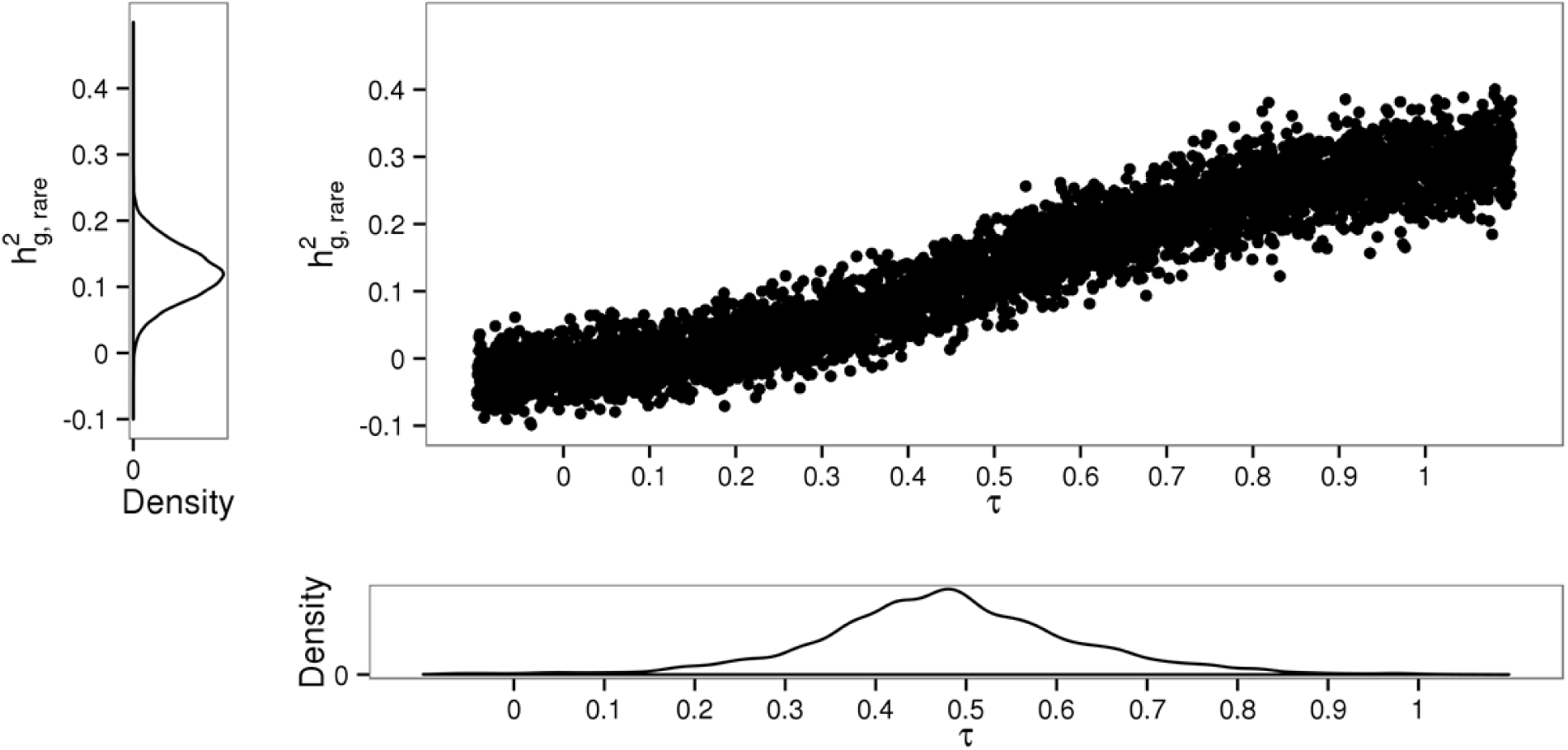
Relationship between strength of selection, coupling parameter τ, and allelic effect sizes in PrCa using heritability partitioning for the African ancestry sample. Figure a) shows the density estimate for 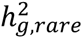 obtained from real data. Figure b) depicts the influence of τ on 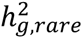. Each point represents an estimate of 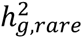 given phenotypes simulated from real genotype under the Eyre-Walker model. Figure c) displays the estimated empirical density of τ. Estimates were obtained by matching a sampled value of 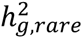 from Figure a) with the closest point-estimate from Figure b).

We produced a total of 9,237 next-generation Illumina sequencing libraries from four ethnic groups (4,006 African, 1,753 European, 1,770 Japanese and 1,708 Latino) using a high throughput library construction strategy that we previously described in ref^33^(see Methods). Results of the sequencing are presented in Table 1, where the mean coverage and the total number of variants discovered is provided for each ancestry group. The total number of mega-bases targeted, the mean coverage, the number of sites discovered, and other metrics for each region is provided in Supplementary Table 1. The average coverage across samples was 9.3x, with a standard deviation of 2.7 across individuals and 2.7 across targeted nucleotides. Altogether, we found a total of 197,786 variants out of which 44% were not identified in the 1000 Genomes project (see Supplementary Table 2). The coverage we obtained for the great majority of samples was high enough in theory to obtain reliable diploid genotype calls after imputation at most targeted bases^34^. To assess accuracy of sequencing, we measured the Pearson correlation between the genotyping calls made with arrays (roughly half of the samples also had been assayed using GWAS arrays). The correlation between sequencing and arrays was r^2^ > 0.82 for common SNPs across all populations with lower correlation towards rare variants (as expected, see Supplementary Figure 2).

**Table 1.** Sizes for each cohort and the coverage and standard deviations in coverage achieved.

| Ancestry | Number of Samples |  | Average coverage per sample (std) | Average coverage per locus (std) | Variants |  |  |
| --- | --- | --- | --- | --- | --- | --- | --- |
|  | Cases | Controls |  |  | Rare | Common | Total |
| | | | | | $0.1\% \leq \text{MAF} < 1\%$ | $\text{MAF} \geq 1\%$ | |
| African | 2,054 | 1,952 | 8.3 (5.1) | 8.4 (5.2) | 58,699 | 63,972 | 122,671 |
| European | 900 | 853 | 8.8 (6.0) | 8.8 (6.0) | 33,606 | 53,164 | 86,770 |
| Japanese | 914 | 856 | 11.8 (5.2) | 11.9 (5.2) | 29,121 | 40,742 | 69,863 |
| Latino | 864 | 844 | 8.0 (5.7) | 8.1 (5.7) | 46,374 | 45,932 | 92,306 |
| Overall | 4,732 | 4,505 | 9.3 (2.7) | 9.4 (2.7) | - | - | - |

**Table 2.**
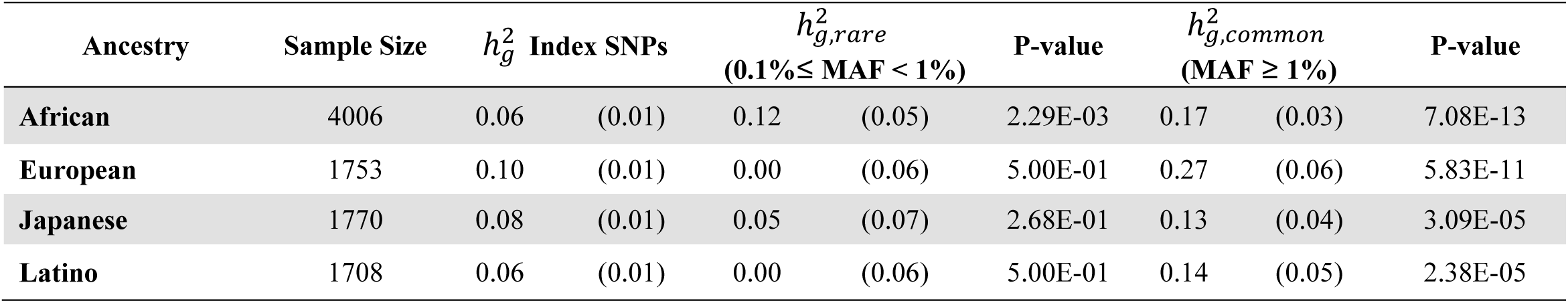
Estimates of 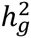 (standard errors) using sequencing data. The results for index SNPs correspond to 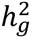 contributed solely from the targeted index variants. Estimates for 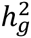 attributable to rare and common components were obtained from joint REML analysis on the underlying liability scale.

### Sequencing explains additional variance beyond the GWAS associated SNPs

To explore the value of sequencing in explaining additional variance in PrCa risk, we fit the genetic data to variance components models to estimate the contribution of all genetic variants at the sequenced risk loci to the underlying liability of PrCa. First, we used simulations starting from the real genotype data to quantify potential biases in variance components estimation. Consistent with findings of previous studies^35^, our simulations show that the approach of utilizing two variance components, one for the rare variants (0.1%≤MAF<1%) and one for common variants (MAF ≥1%), estimated from dosage data and fitted jointly using REML as implemented in GCTA^36^ produces the least amount of bias when estimating SNP-heritability (see Supplementary Table 40; Supplementary Figures 3-10). We also investigated the performance of AI-REML versus EM-REML as implemented in GCTA with AI-REML attaining the least amount of bias in our data (see Supplementary Note 1; Supplementary Tables 17-18; Supplementary Figure 17). We considered the effect of estimating SNP-heritability from best-guess calls rather than imputed dosages and found that both approaches give statistically indistinguishable results. Lastly, we explored the role of linkage-disequilibrium adjustment in estimating the GRM and observed upward bias for LD adjusted GRMs when the underlying 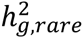 was set to 0 in our simulations. This upward bias was also reflected on estimates using real phenotype data (see Supplementary Table 38). Similar results were obtained over a variety of simulated disease architectures with various amounts of rare variation contribution and total number of underlying causal variants (see Supplementary Note 1).

Motivated by our simulation findings, we estimated the contribution of rare and common variation to risk of prostate cancer by fitting two variance components in GCTA while correcting for the top 10 PCs and age; we report heritability estimates on the liability scale (see Methods; see Supplementary Figure 16 for PCA plot). We find that the total variance explained by all variants at these loci (SNP-heritability 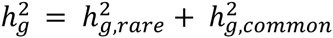) is significantly larger that what is explained by the index variants alone (see Table 2). For example, we estimate the variance explained by all variants in the African ancestry sample at 0.30 (s.e. 0.06), which is significantly larger than the variance explained by all the 84 index variants present in these data (0.05, s.e. 0.01) (6 of the targeted 90 SNPs were not covered by reads passing our analysis filters). This finding is consistent across all ethnicities, thus emphasizing the utility of sequencing in recovering additional signal beyond the index GWAS variants^37^.

Next, we searched for genetic heterogeneity by ancestry in prostate cancer risk using a bivariate REML analysis^38^. Briefly, we computed a single GRM for each unique pair of ancestry groups over the set of SNPs common to both ancestry groups (see Methods) and estimated the genetic correlation using the software GCTA^36^. We then tested the hypotheses that there is no shared genetic liability (SNP-rg=0) or that liability is completely shared (SNP-rg=1) (see Methods). We find significant heterogeneity (after accounting for the 6 pairs tested) for the African and European ancestries (SNP-rg=0.56, s.e. 0.15; p-value_SNP-rg=1_=2.42E-03; see Table 3) and only nominally significant (p-value=0.04) for the Latino and African groups (see Table 3).

**Table 3.** Bivariate REML analysis for each pair of ancestry groups. Estimates of 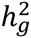 describe the SNP-heritability for each ancestry group over a set of SNPs in common for each pair. Estimates of shared genetic variation in tagged SNPs, or SNP-correlation (SNP-rg). The last two columns give the p-value for the model under the null hypotheses that no correlation exists (SNP-rg =0) and that perfect correlation is present (SNP-rg=1).

| Ancestry pair | Sample Sizes | # SNPs in common | $h_g^2$ (SE) | | Covariance | SNP-rg (SE) | | P-value (SNP-rg = 0) | P-value (SNP-rg = 1) |
| --- | --- | --- | --- | --- | --- | --- | --- | --- | --- |
| African | 4006 | 46332 | 0.20 | (0.03) | 0.17 | 0.56 | (0.15) | 2.10E-04 | 2.42E-03 |
| European | 1753 |  | 0.25 | (0.05) |  |  |  |  |  |
| African | 4006 | 31954 | 0.18 | (0.03) | 0.13 | 0.99 | (0.16) | 9.86E-08 | 0.48 |
| Japanese | 1770 |  | 0.13 | (0.03) |  |  |  |  |  |
| African | 4006 | 61894 | 0.21 | (0.03) | 0.10 | 0.61 | (0.20) | 7.65E-04 | 0.04 |
| Latino | 1708 |  | 0.15 | (0.05) |  |  |  |  |  |
| European | 1753 | 37871 | 0.26 | (0.05) | 0.15 | 0.88 | (0.18) | 3.18E-06 | 0.27 |
| Japanese | 1770 |  | 0.13 | (0.04) |  |  |  |  |  |
| European | 1753 | 58708 | 0.27 | (0.06) | 0.22 | 0.94 | (0.19) | 2.56E-07 | 0.38 |
| Latino | 1708 |  | 0.18 | (0.05) |  |  |  |  |  |
| Japanese | 1770 | 39485 | 0.13 | (0.04) | 0.11 | 1.00 | (0.20) | 3.12E-06 | 0.50 |
| Latino | 1708 |  | 0.18 | (0.04) |  |  |  |  |  |

Having established evidence of heterogeneity, we quantified the contribution to SNP-heritability across the minor allele frequency spectrum in each ancestry group independently. Rare variants explained a significant amount of SNP heritability 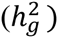 in African ancestry individuals 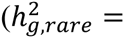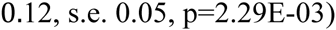, and indeed the proportion of heritability explained by these rare variants is comparable to the heritability explained by common variants at these loci 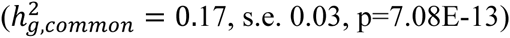. We did not observe significant contribution of rare variation to heritability in the other ethnic groups, although given the limited sample sizes for the other groups, we could not exclude the possibility that the fraction of prostate cancer heritability attributable to rare variants was the same in the other groups. In most of the analyses of heritability stratified by frequency that follow, we focus on people of African ancestry, where we have the highest power to carry out these studies.

We investigated whether the large contribution from rare variants in men of African ancestry is an artifact of data quality (see Supplementary Note 1). We estimated 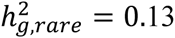 (s.e. 0.06) for the African American ancestry group after removal of any SNPs whose missing-ness before imputation was associated with the trait (p≤0.01) (see Supplementary Table 19). Similar results were obtained when estimating SNP-heritability directly from the hard-calls before imputation both with and without the differentially missing SNPs for the African American group (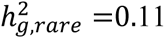, s.e. 0.05; see Supplementary Table 20). To quantify whether hidden relatedness impacts our results, we estimated heritability at various relatedness thresholds; this did not significantly impact the SNP-heritability coming from the rare variants (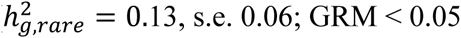; see Supplementary Table 19, see Supplementary Figure 18 and Supplementary Table 41 for distribution of pairwise relatedness values; see Supplementary Tables 21-28 for other ancestry groups). We also explored the role of sequencing coverage and estimated SNP-heritability from GRMs computed after removing SNPs at various levels of coverage. Overall, we found no significant decrease in 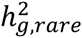 until a large fraction of the SNPs were discarded (coverage ≥7x; see Supplementary Table 43). To rule out potential tagging of signal from other loci in the genome, we repeated the SNP-heritability estimation including a third variance component that includes genotype calls from arrays from the rest of the genome; this yielded similar results for 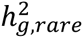 (see Supplementary Tables 29.30; Supplementary Tables 31-36 for other ancestry groups). To account for possible confounding from population substructure, we re-estimated the variance attributable to the rare-frequency class in the African ancestry sample separating Ugandan and non-Ugandan ancestry as well as the 8q24 locus which is known to have a high contribution to risk of PrCa. Overall we find no significant change in 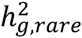 for the African American subset with and without the 8q24 region (see Supplementary Table 4). We also considered bias in our initial estimates of 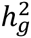 resulting from potential misspecification of the genetic relationship matrix (GRM). Specifically, we estimated variance components using non-standardized genotype data^35^ (which reduces the impact of rare variants in the GRM computation) and found a similar contribution from the rare variants spectrum (see Methods). We also standardized the GRM based on the expected variance rather than the sample estimate and found no significant change 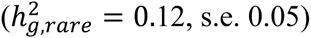. We investigated potential bias in GCTA estimates from linkage across variants of various frequencies by repeating the analysis with 3 variance components corresponding to rare (0.1%≤MAF<1%), low-frequency (1%≤MAF<5%), and common (MAF≥5%) variants; we observed no significant difference in the amount of variance attributable to the rare variant class (Supplementary Table 9). As the standard errors reported by GCTA are asymptotic, we employed a leave-one-out Jack-knife to estimate 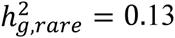 with s.e=0.06 in the African ancestry group (Supplementary Table 37). To further investigate the significance of 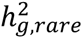in African data, we estimated 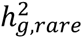 in 1000 simulated phenotypes starting from the real dosage data where the true *h*^2^_*g.rare*_ was set to 0 (i.e. all causal variants were set to have MAF ≥ 1%). In none of the 1000 runs did we observe an estimated 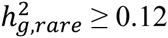, giving an empirical p-value <1/1000. Finally, we performed variance-components analyses using genotypes obtained from best-guess calls, as well as standard unconstrained REML analyses. Overall we found that most of these potential sources of bias are unlikely to significantly change our results (Supplementary Tables 5-8, Supplementary Figures 3-10, see Supplementary Note 1).

**Table 4.** Estimates of τ for each ancestry group under our simulation-based pipeline with MAF partitioning at 1%. Meta-analysis results were computed using an inverse-weighted variance approach. Similar results were obtained with a 5% partitioning on MAF (see Supplementary Figure 11).

| Ancestry | Sample Size | Mean $\tau$ | 95% Confidence Interval | | $h^2_{g,rare}$ | |
| --- | --- | --- | --- | --- | --- | --- |
| African | 4006 | 0.48 | 0.19 | 0.78 | 0.12 | (0.05) |
| European | 1753 | 0.28 | -0.08 | 0.90 | 0.00 | (0.06) |
| Japanese | 1770 | 0.38 | -0.07 | 0.92 | 0.05 | (0.07) |
| Latino | 1708 | 0.39 | -0.08 | 1.05 | 0.00 | (0.06) |
| Meta-analysis | 9,237 | 0.42 | 0.22 | 0.62 | 0.05 | (0.03) |

### Variance partitioning yields insights into the coupling of selection and allelic effect size

In the case of neutral genetic variation, alleles that have a MAF<1% account for only a few percent of genetic variation in the population. However, our empirical results from this study show that at loci known to harbor common variants conferring risk for prostate cancer, variants with MAF<1% account for an order of magnitude larger heritability for the disease. The only plausible explanation for this observation is that newly arising mutations that confer risk for prostate cancer—especially mutations of strong effect—are often subject to strong selection, which prevents them from ever becoming too common.

To quantify the extent to which selection is driving down the frequency of alleles that confer risk for prostate cancer, we derived a simulation-based pipeline that uses estimates of 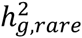 to constrain the value of a parameter τ that Eyre-Walker proposed to measure the coupling between selection coefficients and allelic effect sizes^39^ (see Methods). Briefly, starting from the real genotype data we simulated phenotypes under Eyre-Walker’s model at various values of τ and estimated 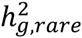 in the simulated trait. We then compared the observed heritability in the real data to the simulations while accounting for sampling noise (see Methods). We estimated τ =0.48 with a 95% confidence interval of [0.19, 0.78] for the African ancestry sample under our mapping procedure (see Figure 1). Similar results were obtained using a 5% MAF cutoff in assigning variants to the rare versus common class (see Supplementary Figure 11). We found that our procedure was relatively robust to changes in parameters. For example, when adjusting the effective population size for African ancestry to 7,500 we re-estimated τ =0.46 with a 95% confidence interval of [0.21, 0.78] (see Supplementary Table 39). Although the small contribution from rare variants joint with small sample sizes for European, Japanese, and Latino data sets prohibit us from estimating a tight confidence interval for the Eyre-Walker’s τ in those populations, the results were roughly consistent across populations (Supplementary Figure 22, Table 4). For example, the estimated mean value of τ for the Japanese cohort is 0.38 with a 95% confidence interval of [-0.07, 0.32]. In a meta-analysis over all ancestry groups we estimated τ=0.42 [0.22, 0.62], which is similar to the African ancestry estimates (unsurprisingly, since the African ancestry data contributes the most to this analysis).

### Single Variant Association

An advantage of full sequencing data—even with a 10-times lower sample size than the largest current GWAS—is that it interrogates all variants in the analyzed samples, and thus has the potential to detect causal variants that are not genotyped/imputed in GWAS studies. We performed marginal association testing at all sequenced variants (n=197,786), and replicated most of the GWAS loci (Supplementary Tables 10-15). We observed a marginal increase in the association signal when including rare variants 0.1%≤MAF<1% across all populations as evidenced by a decrease in the top −log10 p-value (Supplementary Tables 10-15) and a slight enrichment of low p-values in a burden test (Supplementary Figure 24). However, a limitation of the present study is its modest sample size compared to the sample size of 87,040 individuals in the most recent GWAS meta-analyses^24^. For example, of the 84 recovered index variants (6 of the targeted 90 SNPs were not covered by reads passing our analysis filters), only 7 had p-value<10^−8^ (most at 8q24) and 13 had a p-value<10^−4^. Intuitively, this makes sense. Even though we can directly access alleles not on SNP arrays through our targeted sequencing, the advantage we obtain by directly genotyping the SNPs is more than counterbalanced by the 10-fold larger GWAS meta-analysis that has conducted imputation for fine-mapping of common alleles at these regions. To explore additional signal beyond the known index variants, we performed a conditioning analysis (see Methods) on the index variants and observed no variants with p-value<10^−8^ after conditioning; QQ-plots showed residual signal only in the African ancestry sample, consistent with the hypothesis that there is an additional signal beyond the known variants (either due to better tagging of a single causal variant or multiple causal alleles^37^) at these loci (see Supplementary Figures 19-20).

To investigate sequenced SNPs as plausible causal alleles, we integrated epigenetic and genetic data using PAINTOR^40^ to estimate posterior probabilities for causality at each SNP. We used the meta-analysis results for SNPs with MAF ≥ 1.0% (as the Wald statistic is unreliable at MAF<1% and therefore not well-suited to estimation within the PAINTOR framework). First, we ran PAINTOR independently for each of the 20 functional categories that have previously been implicated in PrCa^41^ and found a significant enrichment for causal variants in FOXA1 sites assayed in LNCaP cell lines as well as in the Androgen Receptor binding sites^41^ (Supplementary Figure 23). Second, we selected the functional categories with significant enrichment (at a nominal level of p-value ≤ 0.05) for a joint PAINTOR model to estimate posterior probabilities for each SNP to be causal. Of the 24,840 common variants found in all ethnicities, we observed 9 variants with PAINTOR posterior probability > 0.90 to be causal. In particular, two variants (rs78416326 and rs10486567) exhibited values > 0.99 due to a combination of strong association signal and overlap with functional elements (see Supplementary Table 16, Supplementary Figure 12). Although biological causality cannot be proven on the basis of statistical association alone, we highlight the variants with high posterior probability for follow-up validation.

## Discussion

We have used large-scale targeted sequencing to study the contribution of rare variants to the heritability of PrCa for individuals of diverse ancestry. We find that the total variance in the trait contributed from these regions is significantly greater than the variance localized to known GWAS variants, thus showing that large-scale sequencing can uncover missing heritability. We also provide evidence of heterogeneity by ancestry as well as the first direct evidence of which we are aware of rare variants contributing a substantial fraction of the genetic heritability for a common disease in people of African ancestry. On first principles, there are reasons to think that our results actually underestimate the fraction of heritability due to rare variants. Firstly, our study does not have a sample size sufficient to interrogate extremely rare variants (frequency << 0.1%). Secondly, we focused on known GWAS regions that were ascertained based on harboring an association to a common variant, thus guaranteeing that common variants will be responsible for a substantial fraction of prostate cancer risk at these locations. Prostate cancer is a late-onset disease that primarily affects people after reproductive age. For diseases with younger onset it is plausible that the coupling of selection to disease risk could be even higher; the situation for other diseases can be investigated by large sample size sequencing studies similar to this one. These results motivate additional large sequencing efforts in diverse populations to fully explore the abundance of rare variants that might contribute a substantial fraction of the heritability for at least some important human phenotypes.

We conclude with several caveats of our study. Although we captured the majority of variants at the risk regions through sequencing, a small fraction of the variation is not interrogated in this study due to technological constraints or limited coverage. Second, assaying SNP heritability using variance components makes inherent assumptions often invalidated in empirical data; although we ruled out most types of biases estimation through simulations and real data analysis, other biases could still occur.

## Methods

### Datasets

#### The Multiethnic Cohort.

The Multiethnic Cohort (MEC) consists of over 215,000 men and women enrolled from Hawaii and the Los Angeles region between 1993 and 1996^42^. Participants are primarily of Native Hawaiian, Japanese, European American, African American, or Latino ancestry, and were between the ages of 45 and 75 at baseline when they completed a detailed questionnaire to collect information on demographics and lifestyle factors, including diet and medical conditions. Over 65,000 blood samples were collected from study participants for genetic analysis between 1995 and 1996. To obtain information on cancer status, stage and severity of disease, MEC participants were referenced against population-based Surveillance, Epidemiology and End Results (SEER) registries in California and Hawaii. Unaffected cohort participants with blood samples were selected as controls (for case-control sample sizes see Table 1; for stage and grade of cases see Supplementary Table 44).

#### Uganda Prostate Cancer Study (UGPCS).

UGPCS is a case-control study of prostate cancer in Kampala Uganda that was initiated in 2011. Patients diagnosed with prostate cancer were enrolled from the Urology unit at Mulago Hospital, while undiagnosed men (i.e., controls) were enrolled from other clinics (e.g., surgery) within the hospital. All consenting patients who satisfied strict inclusion criteria (cases: > 39 years of age; controls: > 39 years of age, PSA level < 4 ng/ml to dismiss possible undiagnosed prostate cancer) were recruited into the study. Written consent was obtained and two identical informed-consent forms translated into Luganda were provided to each participant for them to read or to be read to them, sign or thumb print. Descriptive and prostate cancer risk-factor information was collected from interviews conducted with patients using a standardized questionnaire. Biospecimens were collected using Oragene saliva collection kits.

### Library preparation and target enrichment

We prepared NGS libraries from all DNA samples following a cost-effective library preparation protocol developed for this study, which makes it possible to perform multiplexed hybridization enrichment^33^. DNA samples from cases and controls were randomly distributed over 96-well plates to avoid plate effects confounding the results. Each sample was molecularly barcoded during the library preparation stage in 96-well plates to allows us to pool many samples for hybrid capture enrichment and subsequent sequencing. We pooled typically 24 samples in equimolar ratio per capture reaction using the custom SureSelect capture reagent described above. In short, we defined the target region to consist of linkage disequilibrium blocks surrounding all PrCa risk variants known at the time of design (October 2011), all coding sequences surrounding the variants within a 200kb window on either side, and evolutionary conserved elements defined by a 29 mammal alignment^31^. This resulted in a total target size of 16.7Mb, of which probes could be designed for 12Mb. The missing 4.7Mb were non-unique regions of the genome that were filtered out according to Agilent design recommendations. An overview table of targeted genes, the variants, the size of the targeted region per variant, and the size of the baited region is given in Supplementary Note 2. Sequencing was performed at Illumina (San Diego, CA, USA) using HiSeq 2000 instruments for 100 cycles paired end sequencing. Using this approach we covered 78% of the targeted regions (see Supplementary Table 1) of which, 26 regions (41%) expressed mean coverage ≥ 10x.

### Alignment and genotype calling

Sequences were aligned to the human genome reference sequence (*hg*19) using BWA version 0.6.1^43^. Variants were called using the GATK best practices workflow^44^, including mapping the raw reads to the reference genome, base recalibration and compression, and joint calling and variant recalibration. After QC, 11.3Mb of autosomal sequence was considered; due to complexities in the analysis we disregarded data on the X chromosome. Starting from the GATK likelihoods, we applied LD-aware genotype calling using Beagle^45; 46^ version 3.3.2 using the 1000 Genomes v3 data as reference. Variants that displayed low-quality calling (r^2^ < 0.6) or MAF < 0.1% were dropped from the analysis (n= 588,410) resulting in 197,786 SNPs across all ancestry groups. To take advantage of the lower error rate of the GWAS arrays, prior to LD-aware calling, overlapping sequenced SNPs were replaced with their array counterpart. This resulted in 6,028 replaced calls for the African group, 5,395 for the European, 2,642 for the Japanese, and 2,805 for Latinos. The first 10 principal components for each ancestry group were computed using GCTA^36^ from the sequenced common variants (MAF ≥ 1.0%) after LD pruning (r^2^ < 0.2)^47^

### Genotype Array design

To capture SNP-heritability tagged outside of the targeted regions we assayed individuals using the Illumina 1M-Duo for the African ancestry group, Illumina 660W-Quad for Latino and Japanese groups, and Illumina Human610 for Europeans. The number of samples genotyped by array is n=3,078 for the African group, 1,627 for the European, 1,674 for the Japanese, and 1,642 for the Latinos. For quality control we removed any SNP with missing-ness > 0.10. To remove any confounding from tagged variants within the targeted sequenced regions, we removed any SNP within 0.5 Megabases of any region and any SNP with LD > 0.2 of index variants. We further pruned the set to remove any variants with pair-wise LD > 0.3. This resulted in n=251919, 182983, 96711, and 109118 array-based SNPs for the Africans, Europeans, Japanese, and Latinos respectively (see Supplementary Table 21).

### Association analyses

Each variant was subjected to an unconditional marginal case-control association test adjusting for age, Ugandan ancestry for the African group, and the top 10 principal components under a log-additive model performed by PLINK 1.9^48^. All reported p-values are asymptotic estimates from the Wald statistic. We extended the unconditional association test by incorporating the known associated variants (index SNPs) as covariates for each SNP at a given locus. Conditional association tests were implemented in Python 2.7 with the package statsmodels version 0.5. A meta-analysis combining individual population results was performed using METAL^49^ version 2011-03-25. Of the 197,786 analyzed SNPs, 183 were removed from the meta-analysis due to having multi-allelic values when compared across all populations. To perform SKAT-O tests for the African ancestry group, we used a non-overlapping sliding window approach to group rare SNPs into bins containing at most 100 variants across each targeted region resulting in a total of 601 bins. Tests were performed using the software PLINKSEQ version 0.10. To predict the total risk from sequenced variants we performed BLUP prediction in GCTA version 1.24 over a single variance component. Predicted effects were partitioned into rare and common variants and risk-scores computed using the predicted allelic effects with PLINK. Training and prediction was performed using 10-fold cross-validation over samples for each ancestry group (see Supplementary Note 1).

### Heritability analyses

We estimated the genetic relationship matrix (GRM) as 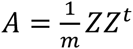, where Z is the standardized genotype matrix and *m* is the number of SNPs. For each sample, two GRMs corresponding to *rare* (0.1% ≤ MAF < 1%) and *common* (MAF ≥ 1%) SNPs were created using GCTA version 1.24. GCTA assumes a linear mixed model where the contribution from each SNP is the result of a random effect given by *y* = *X β* + ∑_*i*_ *g*_*i*_ + ∈ where *y* is a vector of phenotypes, *X* is a covariate matrix (e.g. age), *β* is a vector of fixed effects, and *g*_*i*_ is a vector of random genetic effects for the *i^th^* component (we partition into *g*_*rare*_, *g*_*common*_). The variance of *y* is given by, *var*(*y*) = 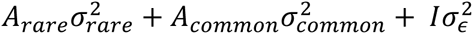, where *A_rare_* and *A_comman_* correspond to the GRMs for rare and common SNPs. Creation of the GRMs was done directly from the dosage data (similar results were obtained using best-guess calls, see Supplementary Tables 6, 7). We estimate the SNP-heritability contributed from rare variants as, 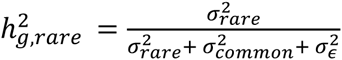. The SNP-heritability analysis was performed on the dichotomous case-control phenotype using constrained REML in GCTA with a prevalence of 0.19 for the African ancestry group, 0.14 for European and Latino, and 0.10 for Japanese (seer.cancer.gov). Hence, all reported values of 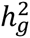 are on the underlying liability scale. To estimate the contribution of the known index variants to SNP-heritability, we computed a GRM restricted to the 84 known variants. The covariate matrix for each ancestry group consisted of age and first 10 principal components (with an additional binary variable indicating Ugandan ancestry for the African ancestry group). LD adjusted GRMs were computed using LDAK^50^ version 4.2. P-values were estimated from a likelihood ratio test by dropping one component and comparing against the reduced model (as implemented in GCTA). To estimate GRMs from array data, we removed any SNP within 0.5 Megabases of the targeted regions and further pruned for pair-wise LD > 0.2 in addition to any remaining variants in LD with index SNPs (see Supplementary Figure 21). For bivariate REML analysis we define the GRM for samples over two ancestry groups as 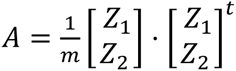 where *Z*_*i*_ is the standardized genotype for ancestry group i and m is the number of SNPs shared between both groups^51^.

### Coupling selection with allelic effect size

We investigated the relationship between selection and marginal effect sizes on PrCa risk using the Eyre-Walker model^39^, that sets allelic effect sizes *β* = (4*N*_*e*_|*s*|)^*τ*^ (1 + *∈*). Here, *N*_*e*_ is the effective population size (set to 10,000 for our analyses^52^), *s* is the selection coefficient of the allele, and *∈* is normally distributed noise (*σ*_*∈*_ = 0.5; varying this parameter does not significantly affect underlying rare/common variation^39^). As τ increases we expect the allelic effects, and thus, the contribution to 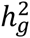 from rare variants, to increase due to rare SNPs experiencing stronger selective pressure than common SNPs (Supplementary Figures 13-15). In order to determine how τ plays a possible role in the underlying architecture for PrCa, we followed a five-step simulation procedure: 1) randomly select a set of 10,000 SNPs to be causal; 2) assign selection coefficients to each causal variant by mapping their allele frequency to selection coefficients^53^; 3) simulate allelic effects under the Eyre-Walker model given selection coefficients, τ and (*σ _∈_* = 0.5; 4) simulate a continuous trait starting from the real genotype data with total SNP-heritability matching the SNP-heritability estimated from real data; 5) perform joint REML analysis in GCTA to estimate rare and common SNP-heritability for the simulated trait. This procedure was repeated for 5,000 values of τ uniformly distributed over the interval [-0.1, 1.1]. To match the observed results in real data to the results from the simulations, we sampled 10,000 values from 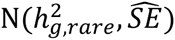 and identified the closest estimate observed under the simulation results and recorded its simulated value for τ. This enabled us to convert the statistical noise around the estimate of the proportion as obtained by GCTA into a variance around τ for each of the ancestry samples.

